# The 5-HT_7_ receptor antagonist SB 269970 ameliorates maternal fluoxetine exposure-induced impairment of synaptic plasticity in the prefrontal cortex of the offspring female mice

**DOI:** 10.1101/2024.02.26.582031

**Authors:** Bartosz Bobula, Magdalena Kusek, Grzegorz Hess

## Abstract

**Background:** Although the use of a selective serotonin reuptake inhibitor fluoxetine in depression during pregnancy is considered safe, it might increase the risk of affective disorders and cognitive symptoms in progeny. In animal models, maternal exposure to fluoxetine throughout gestation and lactation negatively affects the behavior of the offspring. Little is known about the effects of maternal fluoxetine on synaptic transmission and plasticity in the offspring cerebral cortex.

**Methods:** During pregnancy and lactation C57BL/6J mouse dams received fluoxetine (7.5 mg/kg/day) with drinking water. Female offspring mice received intraperitoneal injections of the selective 5-HT_7_ receptor antagonist SB 269970 (2.5 mg/kg) for 7 days. Whole-cell and field potential electrophysiological recordings were performed in the medial prefrontal cortex (mPFC) *ex vivo* brain slices.

**Results:** Perinatal exposure to fluoxetine resulted in decreased field potentials and impaired long-term potentiation (LTP) in layer II/III of the mPFC of female young adult offspring. Neither the intrinsic excitability nor spontaneous excitatory postsynaptic currents were altered in layer II/III mPFC pyramidal neurons. In mPFC slices obtained from fluoxetine-treated mice that were administered SB 269970 both field potentials and LTP magnitude were restored and did not differ from controls.

**Conclusions:** Treatment of fluoxetine-exposed mice with a selective 5-HT_7_ receptor antagonist, SB269970, normalizes synaptic transmission and restores the potential for plasticity in the mPFC of mice exposed *in utero* and postnatally to fluoxetine.

## Introduction

It is generally thought that the use of selective serotonin (5-hydroxytryptamine, 5-HT) reuptake inhibitors (SSRIs) in pregnancy does not increase the risk of major negative outcomes in progeny [1, 2]. However, there are reports demonstrating that prenatal exposure to SSRIs may increase risk of autism spectrum disorder, affective disorders, and cognitive symptoms in later life [3, 4, 5]. Since 5-HT acts as an important regulator of the proliferation, migration, differentiation and establishment of synaptic connections between cortical neurons, maternal SSRIs have the potential to distort the development of the cerebral cortex [6, 7]. One of the most commonly prescribed SSRIs is fluoxetine (FLX) [8, 9].

Experiments on rodents have confirmed that maternal FLX exposure throughout gestation and lactation results in decreased impulsivity, increased depressive-like and aggressive behavior as well as deficits in social communication and interaction, and occurrence of repetitive patterns of behavior in offspring [10, 11, 12]. We have recently reported that FLX-exposed female mice offspring exhibited deteriorated performance in the temporal order memory task and reduced sucrose preference [13, Preprint]. In addition, the offspring of dams exposed to FLX express disturbances in cortical processing of sensory stimuli [14]. These effects are likely to at least partially result from impairment of excitatory synaptic transmission in the medial prefrontal cortex (mPFC). Changes in glutamatergic markers’ levels have been found in the mPFC of animals exposed perinatally to FLX and maternal FLX exposure results in decreased dendritic complexity and spine density of cortical pyramidal neurons in offspring [15, 16, 17]. We have recently reported that maternal FLX exposure throughout gestation and lactation decreased field potentials (FPs), impaired long-term potentiation (LTP) and facilitated long-term depression (LTD) in the prelimbic (PL) subdivision of the mPFC of female but not male offspring mice [13, Preprint]. In the present study we aimed at finding the effects of perinatal FLX exposure on intrinsic excitability of individual pyramidal neurons in the PL cortex of adolescent female mice and their spontaneous excitatory synaptic inputs.

It has been reported that maternal FLX-induced behavioral alterations are reversible by a re-exposure of adult offspring to the drug [12], suggestive of a role of the 5-HT neurotransmission in alleviating the detrimental effects of maternal FLX exposure. We have previously shown that firing of 5-HT neurons of the dorsal raphe nucleus (DRN), that provides 5-HTergic innervation to the forebrain, can be enhanced by administration of the selective 5-HT_7_ receptor (5HT7R) antagonist SB 269970 [18]. We have also shown that treatment of rats with SB 269970 is effective in counteracting repeated restraint stress-induced alterations of glutamatergic transmission and synaptic plasticity in the frontal cortex [19]. Thus, the second aim of this study was to determine whether blockade of 5HT7Rs with SB 269970 could reverse maternal FLX-induced alterations in synaptic transmission and plasticity in the PL cortex of adolescent female mice.

## Materials and methods

### Animals and treatment

The experiments were carried out in accordance with the European Communities Council Directive of September 22, 2010 (2010/63/UE) on the protection of animals used for scientific purposes and national law. Experimental protocols were approved by the 2nd Local Institutional Animal Care and Use Committee at the Maj Institute of Pharmacology, Polish Academy of Sciences in Krakow. C57BL/6J mice (Charles River Laboratories, Sulzfeld, Germany) were kept on a 12/12 h light-dark cycle with light on at 7:00 and off at 19:00. Standard RM3 food (Special Diet Services, Witham, UK) was available *ad libitum*. Mice had free access to tap water before mating and average daily drinking was recorded.

Mice were mated at the age of 75–90 days and females were checked the following day (the gestational day 0, GD 0) for the presence of a vaginal plug. During pregnancy and lactation dams of the experimental group received FLX with drinking water [20] (Fig. 1a). FLX concentration was 0.045 mg/ml for delivering approx. 7.5 mg/kg/day FLX dose based on the weight of dams on GD 0 [10, 21]. To avoid direct FLX exposure in the pups that begin to consume food and water, FLX delivery was discontinued after postnatal day (PND) 18 (Fig. 1a). Control dams had free access to tap water. Offspring of experimental and control dams were weaned on PND 25 and group-housed with free access to standard food and tap water.

**Fig. 1.**
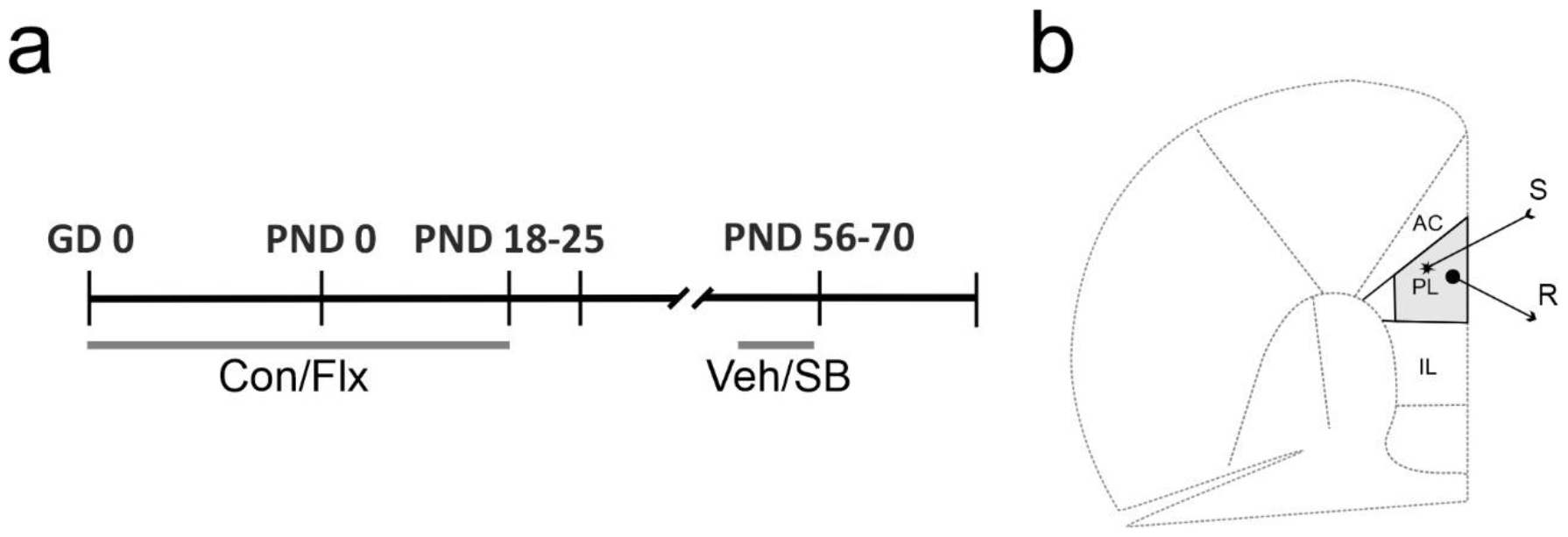
Experimental design. (**a**) Timeline of the procedure. Con/FLX - drinking water alone or fluoxetine in drinking water; Veh/SB - seven daily injections of SB 269970 or its vehicle (0.9% NaCl); GD - gestational day; PND - postnatal day. (**b**) Schematic illustration of a mouse coronal brain slice containing the prefrontal cortex with the recording area marked in grey. AC - anterior cingulate cortex; PL - prelimbic cortex; IL - infralimbic cortex; S - stimulation site; R - recording site. Adapted from [23]

### Whole-cell recording and data analysis

Female FLX-exposed and control offspring mice aged 8–10 weeks (PND 56–70) were anesthetized with isoflurane (Aerrane, Baxter, UK) and coronal slices (300 μm thick) were cut from the frontal cortex using a vibrating microtome (Leica VT1200S) in N-methyl-D-glucamine (NMDG)-based artificial cerebrospinal fluid (ACSF) containing (in mM): NMDG (92), KCl (2.5), CaCl_2_ (0.5), MgSO_4_ (10), NaH_2_PO_4_ (1.2), NaHCO_3_ (30), HEPES (20), D-glucose (25), sodium ascorbate (5), thiourea (2), sodium pyruvate (3), N-Acetyl-L-cysteine (12), pH = 7.3. Slices were then incubated at 35°C for 25 min in while gradually introducing NaCl (the ‘Na spike-in’ method [22]. Slices were stored at room temperature in HEPES-ACSF containing (in mM): NaCl (86), KCl (2.5), CaCl_2_ (2), MgSO_4_ (2), NaH_2_PO_4_ (1.2), NaHCO_3_ (30), HEPES (20), glucose (25), sodium ascorbate (5), thiourea (2), sodium pyruvate (3), N-Acetyl-L-cysteine (12), pH = 7.3, for at least 1 hour.

Individual slices were transferred to the recording chamber and superfused at 2 ml/min with warm (32 ± 0.5 °C), modified ACSF containing (in mM): NaCl (124), KCl (2.5), CaCl_2_ (2), MgSO_4_ (2), NaH_2_PO_4_ (1.5), NaHCO_3_ (24) and D-glucose (12.5), HEPES (5) bubbled with 95% O_2_ – 5% CO_2_. Cells were visualized using Zeiss Axioskop 2 FS upright microscope equipped with a long-range water immersion objective (×40/0.8 NA) and an infrared camera. Recording micropipettes (5–6 MΩ), pulled on P-87 puller (Sutter Instruments, Novato, CA, USA), were filled with a solution containing (in mM) K-gluconate (122), KCl (10), EGTA (0.3), HEPES (10), phosphocreatine (10), Mg-ATP (5) and Na-GTP (0.4); 290 mOsm; pH = 7.3. The liquid junction potential experimentally established for the recording solutions used was approximately 12 mV and the data were not corrected for this offset. Whole-cell recordings were obtained from layer II/III pyramidal cells of the PL. Signals were acquired using the MultiClamp 700B amplifier (Molecular Devices, Sunnyvale, CA, USA) and digitized using the Digidata 1440 interface (Molecular Devices, Sunnyvale, CA, USA) and pClamp 10 software.

Responses to a series of hyper- and depolarizing 500 ms current steps (increment: 20 pA) were recorded in the current-clamp mode. For each cell the relationship between injected current intensity and the number of evoked action potentials was plotted. The gain was defined as a slope of the regression line fitted to experimental data. For recording of sEPSCs, neurons were voltage-clamped at -70 mV for 10 minutes and then recordings were performed for 4 min. Synaptic events were analyzed off-line using the Mini Analysis program (Synaptosoft Inc., ver. 6.0.3). Statistical analysis was carried out using Student’s t-test or Mann-Whitney test (GraphPad Prism 9 software).

### Field potential recording, LTP induction and analysis

For field potential experiments brain slices were prepared from FLX-exposed and control adolescent female offspring that had been subjected to intraperitoneal injections of either SB 269970 (dose: 2.5 mg/kg, volume: 1 ml/kg) or vehicle (0.9% NaCl) once daily, for 7 days (Fig. 1a). Brain slices were prepared 24 h after the last injection to avoid acute effects of the drug.

Coronal slices (380 μm thick) were cut in N-methyl-D-glucamine-based, cold ACSF (see above). A single slice was transferred to the recording chamber of the interface type and superfused with modified ACSF containing (in mM): NaCl (132), NaHCO_3_ (26), KCl (2), CaCl_2_ (2.5), MgSO_4_ (1.3), KH_2_PO_4_ (1.25) and D-glucose (10), bubbled with 95% O_2_ – 5% CO_2_ at 2.5 ml/min (32 ± 0.5 °C). A concentric Pt-Ir stimulating electrode (FHC, Bowdoin, ME, USA) was placed in layer V of the PL cortex (Fig. 1b). Stimuli (duration: 0.2 ms) were applied at 0.016 Hz using a constant-current stimulus isolation unit (WPI). Field potentials (FPs) were recorded using a glass micropipette filled with ACSF (1–2 MΩ), which was placed in cortical layer II/III. FPs were amplified (Axoprobe 1A, Axon Instruments, Foster City, CA, USA), A/D converted at 10 kHz and stored using Micro1401 interface and Signal 2 software (Cambridge Electronic Design, Milton, Cambridge, UK).

A stimulus–response curve was recorded for each slice. To obtain the curve, stimulation intensity was gradually increased stepwise (20 steps; 5–100 μA) and one response was recorded at each stimulation intensity. Next, stimulation intensity was adjusted to evoke a response of 30 % of the maximum FP amplitude and three responses to paired-pulses (50 ms interpulse interval) were recorded and averaged. For the induction of LTP, theta burst stimulation (TBS) was used [24]. TBS consisted of ten trains of stimuli at 5 Hz, each composed of five pulses at 100 Hz, repeated 5 times every 15 s.

The stimulus–response curves were fit with the Boltzmann equation: V_i_ = V_max_/(1 + exp((u − u_h_)/−S), where V_max_ is the maximum FP amplitude; u is the stimulation intensity; u_h_ is the stimulation intensity evoking FP of half-maximum amplitude; S is the factor proportional to the slope of the curve.

The amount of LTP was determined as an average increase in the amplitude of FPs recorded between 45 and 60 min after TBS, relative to baseline. The results are expressed as the means ± SEM. Statistical analysis was carried out using two-way ANOVA followed by Tukey’s *post hoc* test (GraphPad Prism 9 software).

## Results

Measurements of intrinsic excitability of layer II/III pyramidal neurons of the PL cortex did not reveal differences in the gain between pyramidal neurons of FLX-exposed and control mice (0.0311 ± 0.0009 Hz/pA vs. 0.0304 ± 0.0009 Hz/pA, respectively; t = 0.5567, df = 39, *p* = 0.581; t-test; Fig. 2a, b). In cells originating from FLX-exposed and control mice both the resting membrane potential (−72.43 ± 0.87 mV vs. -74.27 ± 0.79 mV, respectively; t = 1.574, df = 39, *p* = 0.123; t-test) and input resistance (226.4 ± 22.12 MΩ vs. 195.4 ± 19.54 MΩ, respectively; t = 1.052, df = 39, *p* = 0.299; t-test) were similar.

**Fig. 2.**
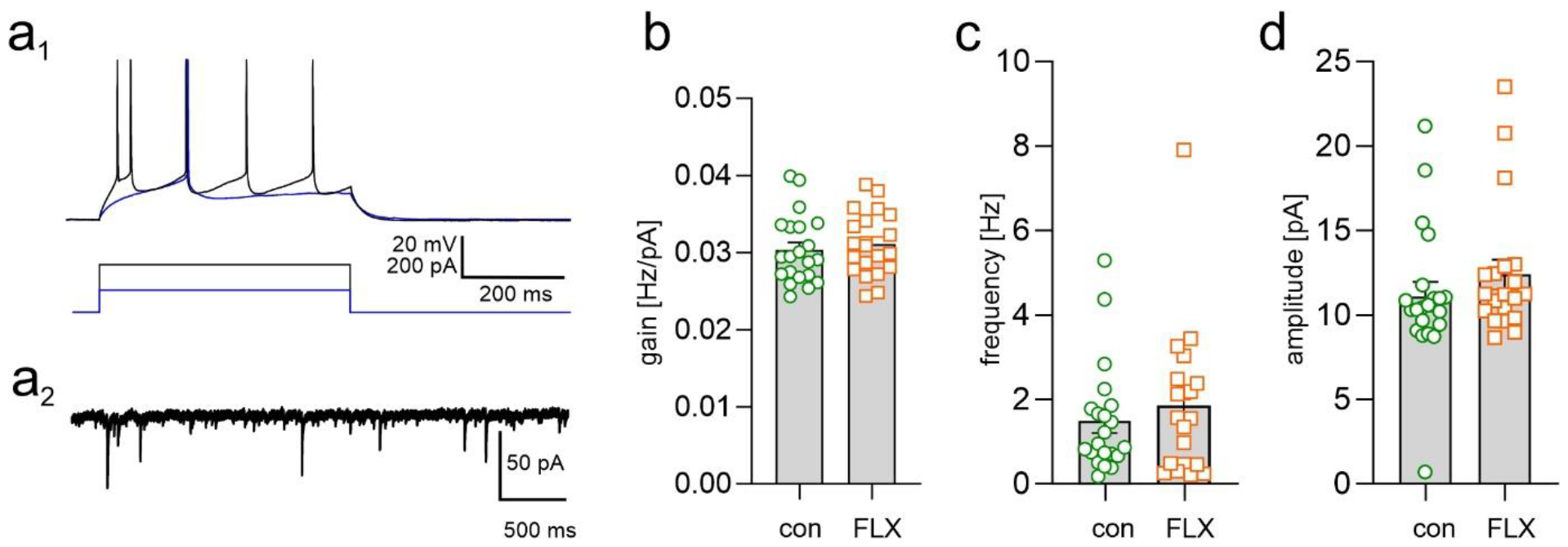
Maternal FLX exposure does not modify the intrinsic excitability of layer II/III mPFC pyramidal neurons and their spontaneous excitatory synaptic inputs. (**a**_**1**_) Superimposed responses of a cell from a control mouse to depolarizing current pulses of two intensities. (**a**_**2**_) An example of a raw sEPSC recording from a control neuron. (**b**) Summary graph of the gain of pyramidal neurons in control (con) and FLX-exposed (FLX) mice. In this and the following graphs mean values ± SEM are depicted with circles and squares representing individual neurons. (**c**) Summary graph showing mean sEPSC frequency. (d) Summary graph showing mean sEPSC amplitude. The differences between groups are not significant.

No FLX-induced changes in were evident in sEPSC frequency (FLX-exposed vs. control: 1.843 ± 0.399 Hz vs. 1.494 ± 0.284 Hz, *p* = 0.670; Mann-Whitney test; Fig. 2c) and sEPSC amplitude (FLX-exposed vs. control: 12.40 ± 0.875 pA vs. 11.09 ± 0.88 pA, *p* = 0.192; Mann-Whitney test; Fig. 2d). In cells originating from FLX-exposed and control mice both the sEPSC rise time (1.747 ± 0.112 ms vs. 1.78 ± 0.128 ms, respectively; t = 1926, df = 39, *p* = 0.848; t-test) and sEPSC decay time constant (5.09 ± 0.27 ms vs. 4.87 ± 0.28, respectively; t = 0.549, df = 39, *p* = 0.585; unpaired t-test) were similar.

To investigate whether the effects of maternal FLX exposure on evoked synaptic transmission and plasticity in the PL cortex could be reversed by treatment with SB 269970, four groups of mice were investigated. Animals that were exposed to FLX and then received either SB 269970 or vehicle (0.9% NaCl) injections for 7 days (Fig. 1a) were termed FLX-SB and FLX-Veh, respectively. Control mice that were not exposed to FLX and received either SB 269970 or vehicle injections were termed Con-SB and Con-Veh, respectively.

Two-way ANOVA of FPs revealed a significant main effect of FLX exposure (F_(1, 48)_ = 14.88, *p* = 0.0003) but no significant main effect of treatment with SB 269970 (F_(1, 48)_ = 1.698, *p* = 0.1987) on the maximum amplitude of FPs calculated using Boltzmann fits (Fig. 3a, b). There was a significant interaction between FLX exposure and treatment with SB 269970 (F_(1, 48)_ = 15.25, *p* = 0.0003). Maximum calculated FP amplitude was significantly lower in the Flx-Veh group, compared to the control (Con-Veh) group (0.95 ± 0.12 mV vs. 1.78 ± 0.13 mV, respectively; *p* < 0.001; Fig. 3a). Treatment with SB 269970 for 7 days resulted in a reversal of the effect of maternal FLX exposure on maximum FP amplitude (Fig. 3a) as it was not significantly different between slices obtained from mice exposed to FLX and then receiving SB 269970 (Flx-SB; 1.51 ± 0.12 mV) and from control (Con-Veh) mice (*p* = 0.2995). Maximum calculated FP amplitude in slices obtained from mice that were not exposed to FLX but were treated with SB 269970 (Con-SB; 1.50 ± 0.12 mV) was also not significantly different from the Con-Veh group (*p* = 0.2718).

**Fig. 3.**
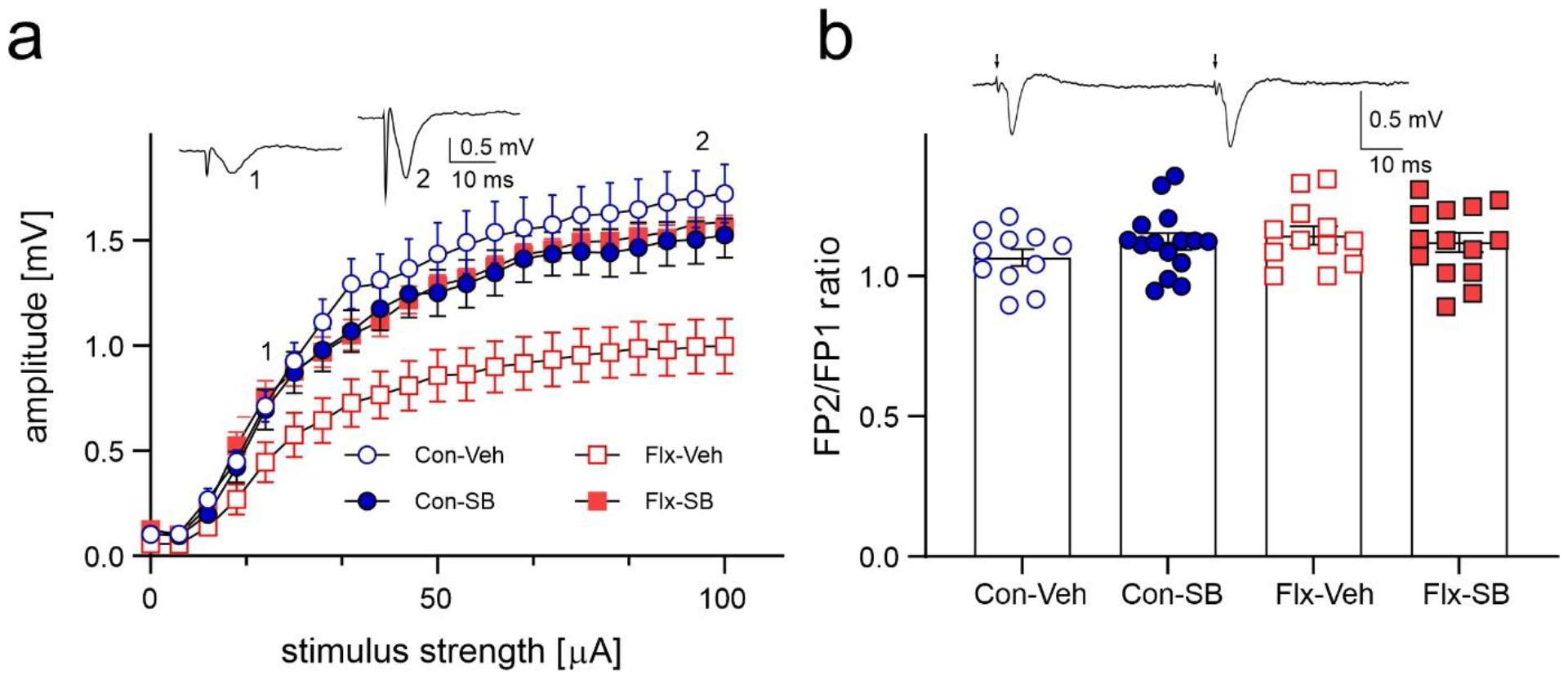
Maternal FLX exposure decreases FP amplitude but does not change paired-pulse response ratio in the mPFC. (**a**) Graph illustrating the effect of FLX exposure and SB 269970 treatment on the relationship between stimulus intensity and mean FP amplitude (± SEM). Red open squares – slices obtained from mice exposed to FLX and receiving vehicle (Flx-Veh); red filled squares – slices from mice exposed to FLX and receiving SB 269970 (Flx-SB); blue open circles - slices obtained from mice not exposed to FLX and receiving vehicle (Con-Veh); blue filled circles - slices obtained from mice not exposed to FLX and receiving SB 269970 (Con-SB). Insets: examples of FPs evoked in a representative Con-Veh slice at two stimulation intensities (1, 2). (**b**) Paired-pulse (FP2/FP1) ratio in four tested groups. Shown are mean values ± SEM with symbols representing individual slices. Inset shows an example of a pair of FPs evoked in a representative Con-Veh slice

Despite a decrease in the maximum FP amplitude due to maternal FLX exposure, ANOVA revealed no main effect of either FLX exposure (F_(1, 48)_ = 1.395, p = 0.2435) or treatment with SB 269970 (F_(1, 48)_ = 0.2791, p = 0.5997) on paired-pulse response ratio of FPs (Fig. 3b). There was also no interaction between these two factors (F_(1, 48)_ = 1.623, p = 0.2097).

Two-way ANOVA of FPs recorded between 45-60 minutes after TBS delivery revealed significant main effects of FLX exposure (F_(1, 48)_ = 14.62, *p* = 0.0004) and SB 269970 treatment (F_(1, 48)_ = 9.727, *p* = 0.0031) as well as their interaction (F_(1, 48)_ = 13.86, p = 0.0005) on the magnitude of LTP (Fig. 4). LTP was smaller in the Flx-Veh group compared to the control (Con-Veh) group (109.7 ± 6.86 % vs. 147.4 ± 5.60 %, respectively; *p* < 0.0001). Treatment with SB 269970 reversed the effect of maternal FLX exposure on LTP as it was not significantly different between slices obtained from mice exposed to FLX and then receiving SB 269970 (Flx-SB; 143.9 ± 8.20 %) and from control (Con-Veh) mice (*p* = 0.9613).

**Fig. 4.**
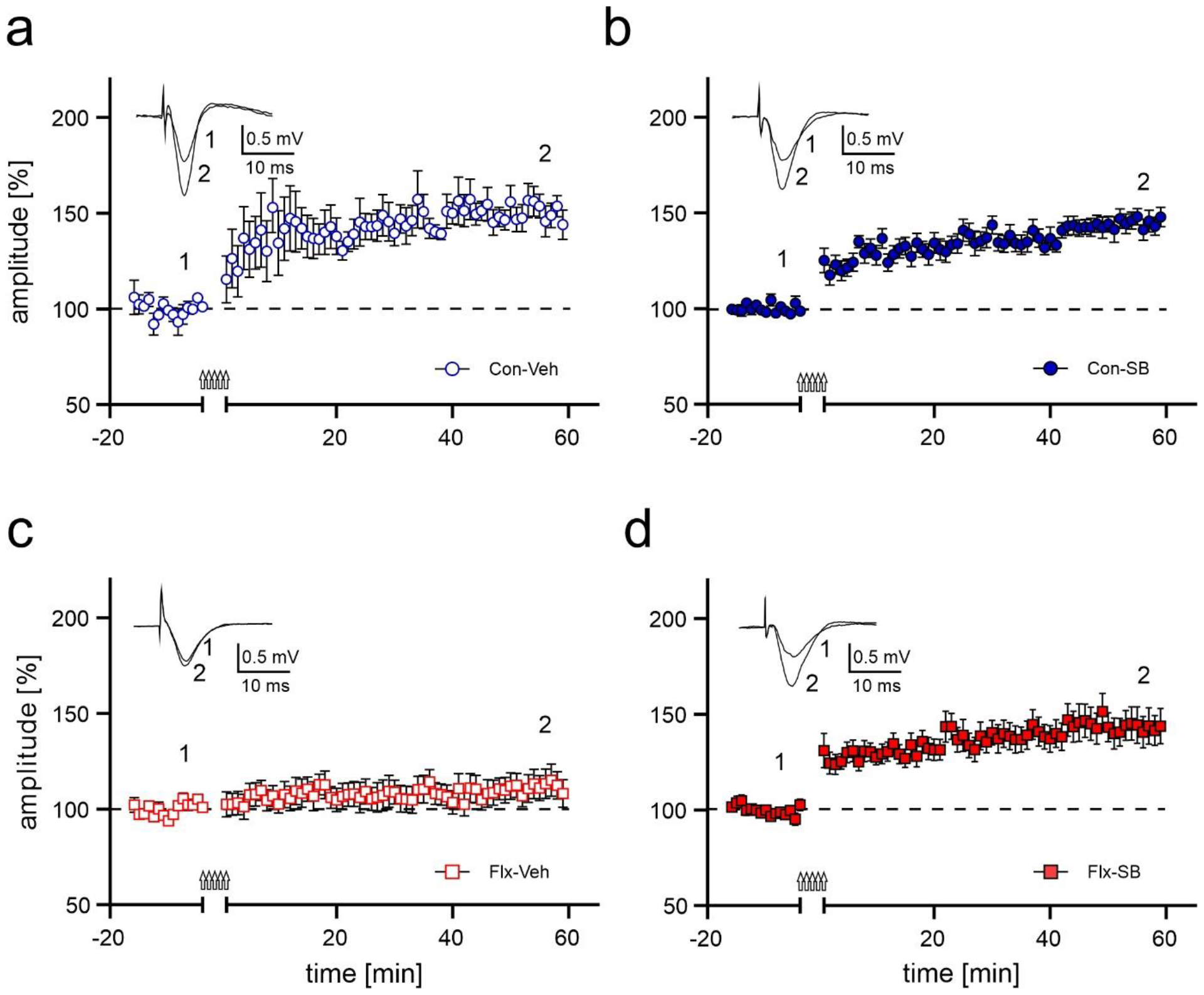
Treatment with SB 269970 reverses the detrimental effect of maternal FLX exposure on LTP in the mPFC. Plots of the time course of FP amplitude (± SEM) before and after LTP induction by TBS (arrows) in slices obtained from (**a**) mice not exposed to FLX and receiving vehicle (Con-Veh), (**b**) mice not exposed to FLX and receiving SB 269970 (Con-SB), (**c**) mice exposed to FLX and receiving vehicle (Flx-Veh) and (**d**) mice exposed to FLX and receiving SB 269970 (Flx-SB). Insets show superpositions of FPs recorded in representative experiments at the times indicated by numbers. Data were calculated as a percentage of changes relative to baseline values

## Discussion

The results of this study confirm that exposure to FLX during pre- and postnatal development decreases the maximum amplitude of evoked FPs as well as LTP magnitude in the PL subdivision of the mPFC of female adolescent offspring mice [13, Preprint]. However, exposure to FLX does not modify the intrinsic excitability and spontaneous excitatory transmission in mPFC pyramidal neurons. Treatment of FLX-exposed mice with a selective 5-HT_7_ receptor antagonist, SB269970, recovers the FP amplitude and restores the potential for long-term synaptic plasticity in the PL cortex.

Short-latency evoked FPs in rodent mPFC slices represent summed, synchronous activity of monosynaptic glutamatergic connections that are activated by presynaptic stimulation [25]. On the other hand, sEPSCs are spontaneous synaptic events that are mostly independent on action potential generation in the presynaptic neuron [26, 27]. Synchronous and spontaneous neurotransmitter release engage different molecular presynaptic mechanisms and, moreover, distinct postsynaptic receptors [28]. Thus, decreased FP amplitude observed in mPFC slices originating from FLX-exposed female mice is consistent with a decreased efficacy of the synchronous mode of the excitatory synaptic activity while the spontaneous mode remains intact.

Disturbances in the functioning of mPFC neural circuitry, including the PL region, have been linked to anxiety- and depression-like behaviors in preclinical models [29]. Proper activity of the PL cortex is important for the ability of rodents to respond adaptively under changing circumstances [30] including active avoidance of aversive events and withholding responses to obtain rewards [31]. The PL cortex contains subpopulations of neurons that are specialized in coding fear memory and its extinction [32]. Importantly, the maintenance of the extinction of learned fear in mice depends on LTP in the mPFC [33]. Thus, impairments of synaptic transmission and LTP that we observed in the PL cortex after maternal FLX exposure are likely to have profound behavioral consequences and are in line with enhanced anxiety observed in rodents exposed to FLX pre- or postnatally [34]. Poor performance of FLX-exposed mice that we observed in the temporal order memory task [13, Preprint] is also consistent with an impairment in functioning of the PL cortex as the temporal order memory depends on the mPFC, and on the PL in particular [35, 36].

The finding that the relationship between amplitudes of paired-pulse evoked responses remained unchanged in FLX-exposed mice, despite generally smaller FPs as compared to control, suggests that the decrease of responses is due to a postsynaptic mechanism, rather than a presynaptic one [37]. However, no changes in the NMDA or AMPA receptor subunit expression were evident in mPFC of Sprague-Dawley rat female offspring after maternal FLX exposure throughout gestation and lactation (10 mg/kg/day) [16]. This discrepancy is likely a result of species difference as the dose of FLX that we used was only slightly lower. Lisboa et al. [10] reported that exposure of mouse dams to FLX during pregnancy and lactation increased the immobility time in the forced swimming test (a depressive-like behavior) in female offspring. We have recently shown that a similar FLX exposure regime decreases sucrose preference, another depressive-like behavior [13, Preprint]. In addition, Millard et al. [16] also reported that maternal FLX exposure (10 mg/kg/day) enhanced forced swim immobility in rat female offspring. Thus, the rat and mouse maternal FLX exposure models are consistent regarding their behavioral effects. It should be noted that a lack of changes in general levels of glutamate receptor subunits in tissue samples after maternal FLX exposure [16] does not exclude the occurrence of functional modifications including changes in receptors’ subcellular distribution and trafficking along with changes in their subunit composition and phosphorylation state [38].

5HT7R blockade has been known to induce antidepressant-like effects that occur faster than those induced by conventional antidepressant drugs [39]. We have previously used SB 269970 to investigate the reversibility of the effects of repeated restraint stress on FPs in the rat frontal cortex. These experiments demonstrated that while the amplitude of layer II/III FPs was increased in stressed animals, LTP magnitude was decreased. However, in slices obtained from rats that had received injections of SB 269970 (1.25 mg/kg) before each of 3 restraint stress session, both FP amplitude and LTP magnitude were not different from controls indicating that repeated stress-induced modifications were prevented by SB 269970 administration [40]. In the rat dorsal raphe nucleus (DRN) prenatal stress increases sEPSC frequency, and repeated administration of SB 269970 (2.5 mg/kg, 7 days) reverses these alterations [41]. We have shown that 5-HT7Rs tonically control firing of inhibitory interneurons that release GABA in the DRN and administration of SB 269970 stimulates the cortical release of 5-HT [42]. Thus, we hypothesize that amelioration of the effects of stress [40, 41] and of maternal exposure to FLX by SB 269970 [13, Preprint; this study] could involve enhanced 5-HT release in target cortical areas.

## Abbreviations

5-HT: 5-hydroxytryptamine, serotonin
5-HT7R: 5-HT_7_ receptor
ACSF: artificial cerebrospinal fluid
DRN: dorsal raphe nucleus
FP: field potential
FLX: fluoxetine
GD: gestational day
LTD: long-term depression
LTP: long-term potentiation
mPFC: medial prefrontal cortex
NMDG: N-methyl-D-glucamine PL prelimbic cortex
PND: postnatal day
sEPSC: spontaneous excitatory postsynaptic current
SSRI: selective serotonin reuptake inhibitor
TBS: theta burst stimulation

## Acknowledgments

The authors thank Marcin Siwiec for improving the English of this article.

## CRediT authorship contribution statement

**Bartosz Bobula:** Methodology, Formal Analysis, Investigation, Visualization, Resources. **Magdalena Kusek:** Formal Analysis, Investigation, Visualization. **Grzegorz Hess:** Conceptualization, Methodology, Formal analysis, Investigation, Writing – Original Draft, Supervision, Funding acquisition

